# Integral methods for automatic quantification of fast-scan-cyclic-voltammetry detected neurotransmitters

**DOI:** 10.1101/2020.04.24.060368

**Authors:** Leonardo X. Espín, Anders J. Asp, James K. Trevathan, Kip A. Ludwig, J. Luis Lujan

**Affiliations:** Department of Neurologic Surgery, Mayo Clinic, Rochester, MN 55905, USA; Mayo Graduate School of Biomedical Sciences, Mayo Clinic, Rochester, MN 55905, USA; Department of Physiology and Biomedical Engineering, Mayo Clinic, Rochester, MN 55905, USA

## Abstract

Modern techniques for estimating basal levels of electroactive neurotransmitters rely on the measurement of oxidative charges. This requires time integration of oxidation currents at certain intervals. Unfortunately, the selection of integration intervals relies on ad-hoc visual identification of peaks on the oxidation currents, which introduces sources of error and precludes the development of automated procedures necessary for analysis and quantification of neurotransmitter levels in large data sets. In an effort to improve charge quantification techniques, here we present novel methods for automatic selection of integration boundaries. Our results show that these methods allow quantification of oxidation and reduction reactions, for multiple analytes, both in vitro and in vivo.

## 1 Introduction

Fast scan cyclic voltammetry (FSCV) is a powerful electrochemical sensing technique that allows quantification of variations in the concentration of electroactive neurochemicals by measuring redox currents resulting from the application of a periodic triangular waveform at a high scan rate [1, 2, 3, 4, 5]. Traditionally, FSCV has depended on the calculation of maximal oxidation currents, measured from known neurochemical concentrations in a solution, which are used to build calibration curves by using linear correlation techniques [6, 7, 8, 9, 10].

Recent studies have exploited the catecholamine adsorption properties of carbon fiber microelectrodes (CFM), to estimate basal concentrations of neurochemicals. [11, 12, 13, 14]. Techniques including fast scan cyclic adsorption voltammetry (FSCAV) use oxidation-charge measurements, rather than maximal currents, which are obtained by time-integrating cyclic voltammograms within intervals containing single oxidation peaks (or “humps”) [11, 12, 13]. However, the accuracy and reproducibility of oxidation-charge measurements are limited by visual selection of integration bounds of the cyclic voltammogram oxidation peaks. In practice, defining which portion of the voltammogram constitutes an oxidation peak (where it begins, and where it ends) is obscured by the noise floor of the dataset, the electrochemical interferents, the presence of artifacts, and background drift. Visual selection leads to ambiguity, introduces additional sources of error and precludes the development of automated procedures necessary for analysis and charge quantification in large data sets.

Here, we describe novel charge quantification techniques by performing automatic selection of integration boundaries. This is achieved by analyzing and identifying voltammogram’s critical, inflection and maximum curvature points, to allow the automatic selection of integration intervals. We test these techniques in both *in vitro* and *in vivo* experimental scenarios.

## 2 Methods

### 2.1 In vitro data collection

In vitro data collection was performed using a FIAlab 3200 flow injection system (FIAlab Instruments, Seattle, WA) and the WINCS Harmoni device [15]. A 110 *µ*m CFM was placed in a flowing stream of artificial cerebrospinal fluid (aCSF) buffer solution with a pH value of 7.4 as described previously [15]. For each measurement, buffered aCSF solutions containing 0.1 *µ*M to 5 *µ*M of either dopamine, adenosine, epinephrine, or norepinephrine (Millipore Sigma, Burlington, MA) were injected for 8 s at 2.25 mL/min.

### 2.2 In vivo data measurements

In vivo measurements were obtained in a rodent model of medial forebrain bundle stimulation and simultaneous FSCV recording in the dorsal striatum as described previously [15]. Rats were sedated prior to surgery with intraperitoneal urethane (1.5 g/kg in a 0.26 g/mL saline solution, Millipore Sigma, Burlington, MA). Analgesia was maintained for rodents with intramuscular buprenorphine (0.06 mg/kg). Both the stimulation electrode and CFM were stereotactically inserted (KOPF instruments, Tujunga, CA). A scalp incision (1.5 - 2.0 cm) was made to expose the skull, and three trephine burr holes (approximately 3 mm in diameter) were drilled to allow implantation of the stimulating, neurochemical sensing, and reference electrodes. Dopamine release was evoked with a series 2 s stimulations using a range of amplitudes from 0.05-2.0 mA and pulsewidths from 0.8 and 1.8 ms, presented in a randomized order. Analyte measurements were obtained by sweeping the CFM potential from a resting potential of −0.4 V to a switching potential of 1.5 V and back to the resting potential, at a rate of 400 V/s every 100ms.

### 2.3 Charge quantification

We define the charge due to a single oxidation/reduction reaction by

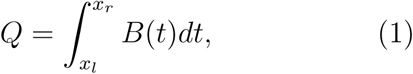

where *B*(*t*) is a background subtracted voltammogram, and we assume that the integration interval [*x*_*l*_ *x*_*r*_] contains a single oxidation/reduction peak (Figure 1a). In this study we use four pairs of integration boundaries (Figure 1) to quantify charge. When there is little or no background drift, indicating a stable background capacitive current, the faradaic current response on a background subtracted voltammogram decays towards zero away of the maximum oxidation current [16]. “True” integration boundaries are defined as the points around an oxidation peak where the current is zero (Figure 1 a). Charges computed with true boundaries provide a useful benchmark for comparing quantification methods. However, “True” limits as defined here are unavoidably confounded by the interplay between the Gaussian dopamine oxidation peak and the noise levels in the recording; as a Gaussian distribution never decays to zero.

**Figure 1:**
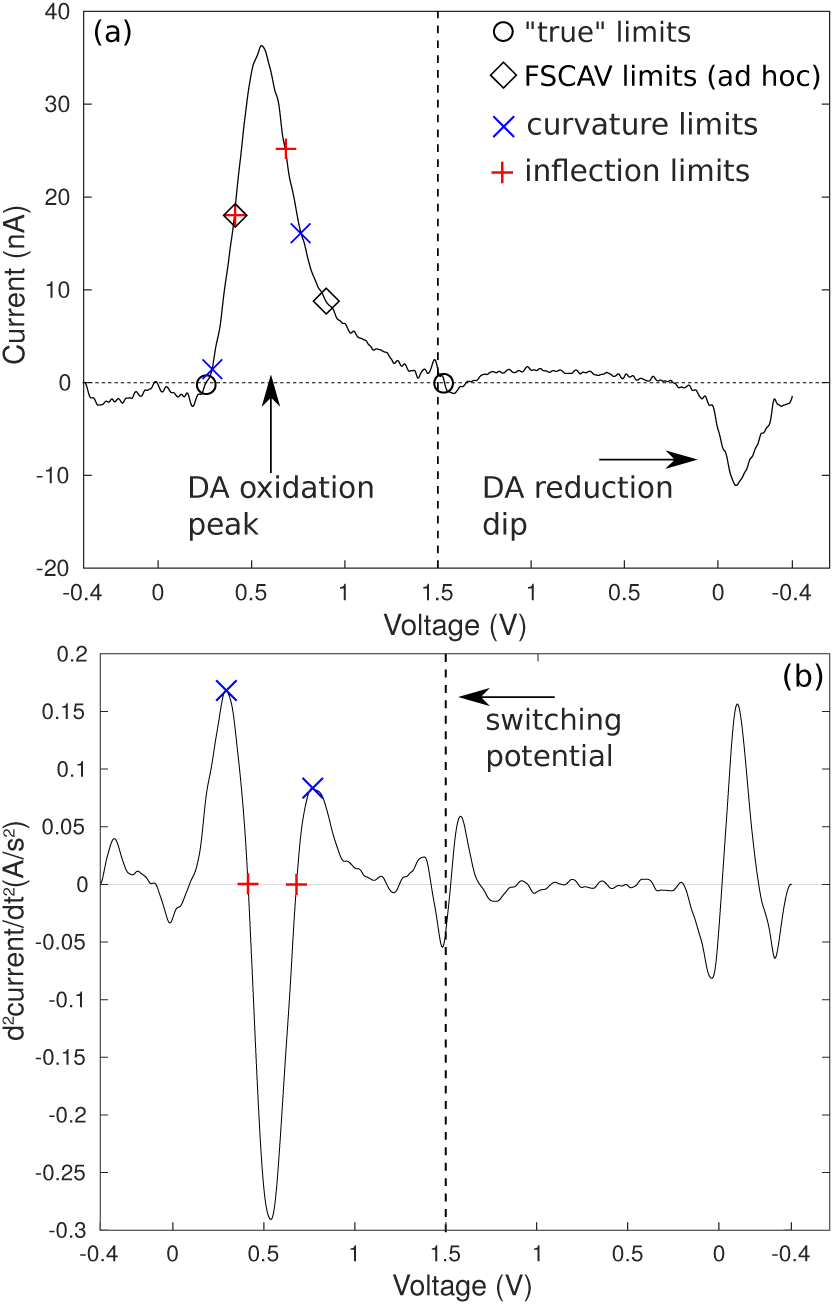
a) Background subtracted voltammogram for a bolus of a 0.5 *µ*M dopamine solution. True (“o”, *x*_*l*_ = 0.26 V and *x*_*r*_ = 1.5 V), FSCAV (“◊”, *x*_*l*_ = 0.4 V and *x*_*r*_ = 0.9 V), inflection (“+”, *x*_*l*_ = 0.41 V and *x*_*r*_ = 0.68 V) and curvature integration boundaries (“x”, *x*_*l*_ = 0.29 V and *x*_*r*_ = 0.76 V) are marked on the voltammogram. b) Second derivative of the voltammogram shown in (a), showing the definition of inflection boundaries (inflection points), and curvature boundaries (the maxima around the location of the oxidation peak).

FSCAV limits (Figure 1 a), correspond to the ad-hoc voltages (0.4 V and 0.9 V) selected as integration limits for the quantification of dopamine [13]. Note that all measurements were collected using a traditional FSCV waveform described in section 2.2 as opposed to FSCAV. Inflection limits are defined as the zeros of the second derivative of a voltammogram, around an oxidation (or reduction) peak, and curvature points (maximum curvature, or maximum second derivative) are defined as the maxima/minima of the second derivative around an oxidation/reduction peak, see Figure 1(b).

Computational routines for filtering and smoothing background-subtracted voltammograms, as well as to calculate higher order derivatives, to find, classify, sort and correct curvature and inflection points are implemented in MATLAB. Correction routines require not adding negative areas to the total charge in the case of oxidation reactions, and have the switching potential as hard limit for *x*_*r*_. Similar considerations are utilized for reduction-charge calculations (excluding regions with positive areas, and using the switching potential as hard limit for *x*_*l*_).

## 3 Results and discussion

### 3.1 Quantification of in vitro catecholamines

In Figure 2 we show (a) true and (b-c) curvature integration boundaries obtained with our algorithms. As a reference we also show the location of the maximum oxidation currents with dots, and panels (b-c) show the FSCAV integration limits taken from reference [11], 0.4 V and 0.9 V with vertical dashed lines. The data for panels (a) and (b) corresponds to background subtracted voltammograms of a flow cell experiment, where 70 dopamine injections with 0.1, 0.5 and 1 micromolar were done. The data for panel (c) corresponds to voltammograms of a flow cell experiment, where 15 norepinephrine injections with 0.1, 0.5, 1 and 5 micromolar were done.

**Figure 2:**
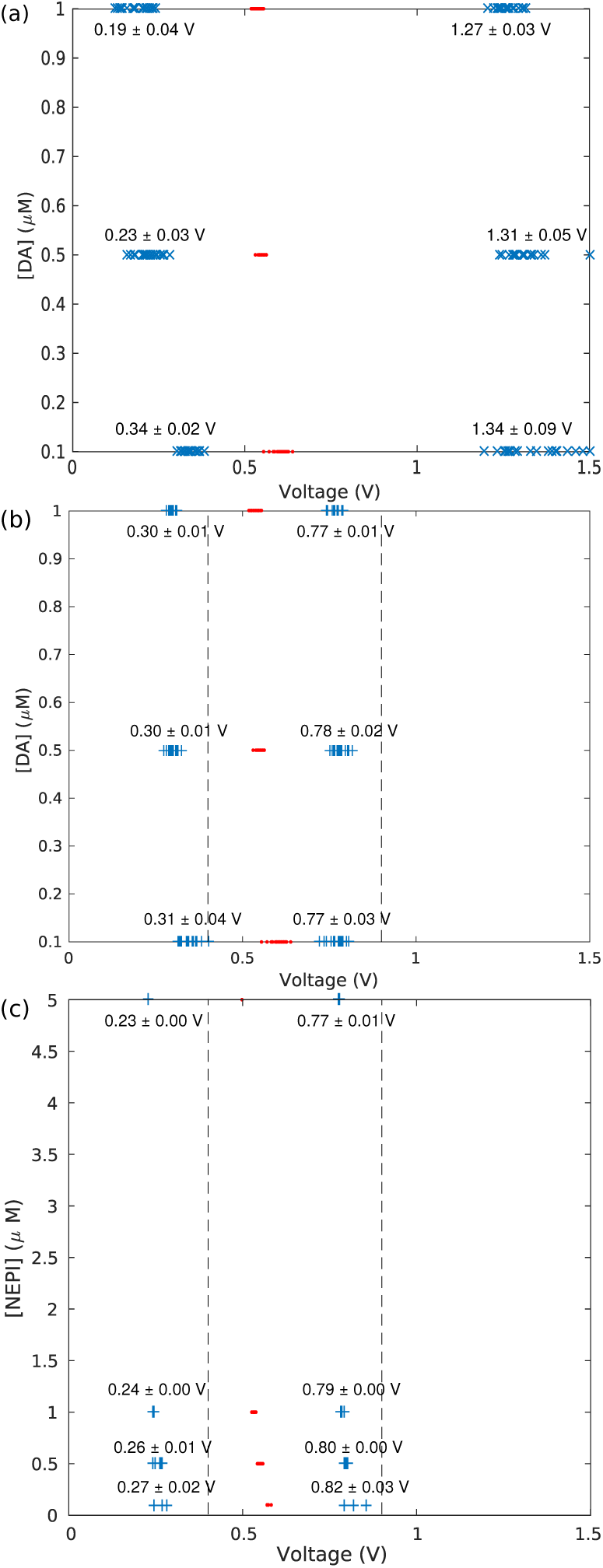
a) True integration limits (“x”) and maximum oxidation currents (dots) obtained from a flow cell experiment consisting of 70 dopamine injections at different concentrations. b) Curvature integration boundaries (“+”) for the experimental data of (a). c) Curvature boundaries for a flow cell experiment with 15 norepinephrine injections at different concentrations. FSCAV limits are shown with dashed vertical lines. Quantities shown in the figure indicate averages and standard deviations.

Panels (a) and (b) of figure 2 show that the curvature boundaries select narrower integration intervals than true boundaries. However, it is interesting to note that curvature boundaries show less variability than the true boundaries (average voltages and standard deviations are shown in Figure 2). This is unexpected given that the computation of curvature boundaries involves calculating higher-order derivatives of voltammograms, which should amplify noise [17], and as a consequence curvature boundaries variability.

Because of the visual similarities between dopamine and norepinephrine background subtracted voltammograms [4, 11, 18], comparisons using norepinephrine data are particularly useful, because they highlight the limitations of using visual identification to obtain integration boundaries. Indeed, panels (b) and (c) of figure 2 indicate that there is a significant difference between average left curvature boundaries for dopamine and norepinephrine voltammograms. As we describe below, this results in curvature boundaries producing more accurate charge quantifications.

Indeed, figure 3 shows oxidation-charge calculations with the four pairs of integration boundaries introduced in section 2.3. Panel (a) shows calculations for the dopamine dataset corresponding to figure 2(a-b), and panel (b) shows calculations for the norepinephrine dataset corresponding to figure 2(c). Figure 3(a) shows that when quantifying dopamine charge, the ad-hoc FSCAV limits and curvature limits show similar accuracy when compared against true charges. However, despite shape similarities between dopamine and norepinephrine background subtracted voltammograms (e. g. correlation coefficient > 0.86, see [19]), Figure 3(b) shows that oxidation charges obtained with curvature limits provide a better approximation to true charges than FSCAV limits. This is consistent with the ad-hoc nature of the FSCAV limits.

**Figure 3:**
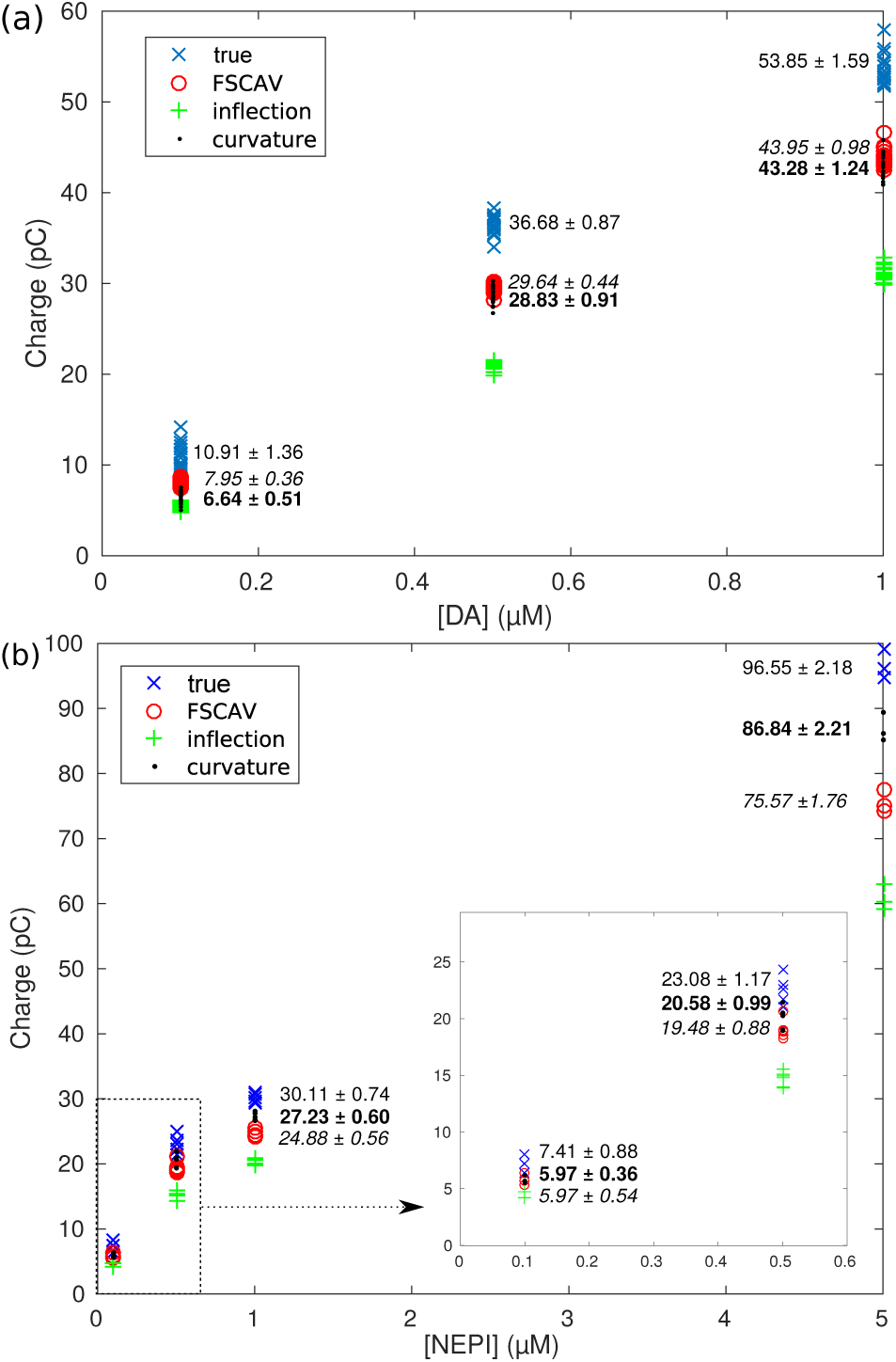
a) Oxidation charges for the data shown in Figure 2(a-b), obtained with different integration boundaries. b) Oxidation charges for the data shown in Figure 2(c), obtained with different integration boundaries. Averages and standard deviations for charges using true, FS-CAV (in italic) and curvature (bold) integration boundaries are also shown.

By design, curvature boundaries select a wider region than that enclosed by the inflection boundaries (figure 1), and they provide a better approximation to the charges obtained by using true boundaries, as seen in both panels of figure 3.

Figure 4 shows an example were curvature integration boundaries provide the best results among the four methods considered. As illustrated by Figure 4(a), epinephrine voltammo-grams have two oxidation peaks [3], and if we try to calculate the oxidation charge due to the primary (or secondary) peak alone, true boundaries produce erroneous results by selecting a region that contains both oxidation peaks. This issue also arises when combinations of analytes (like dopamine and adenosine) are being measured.

**Figure 4:**
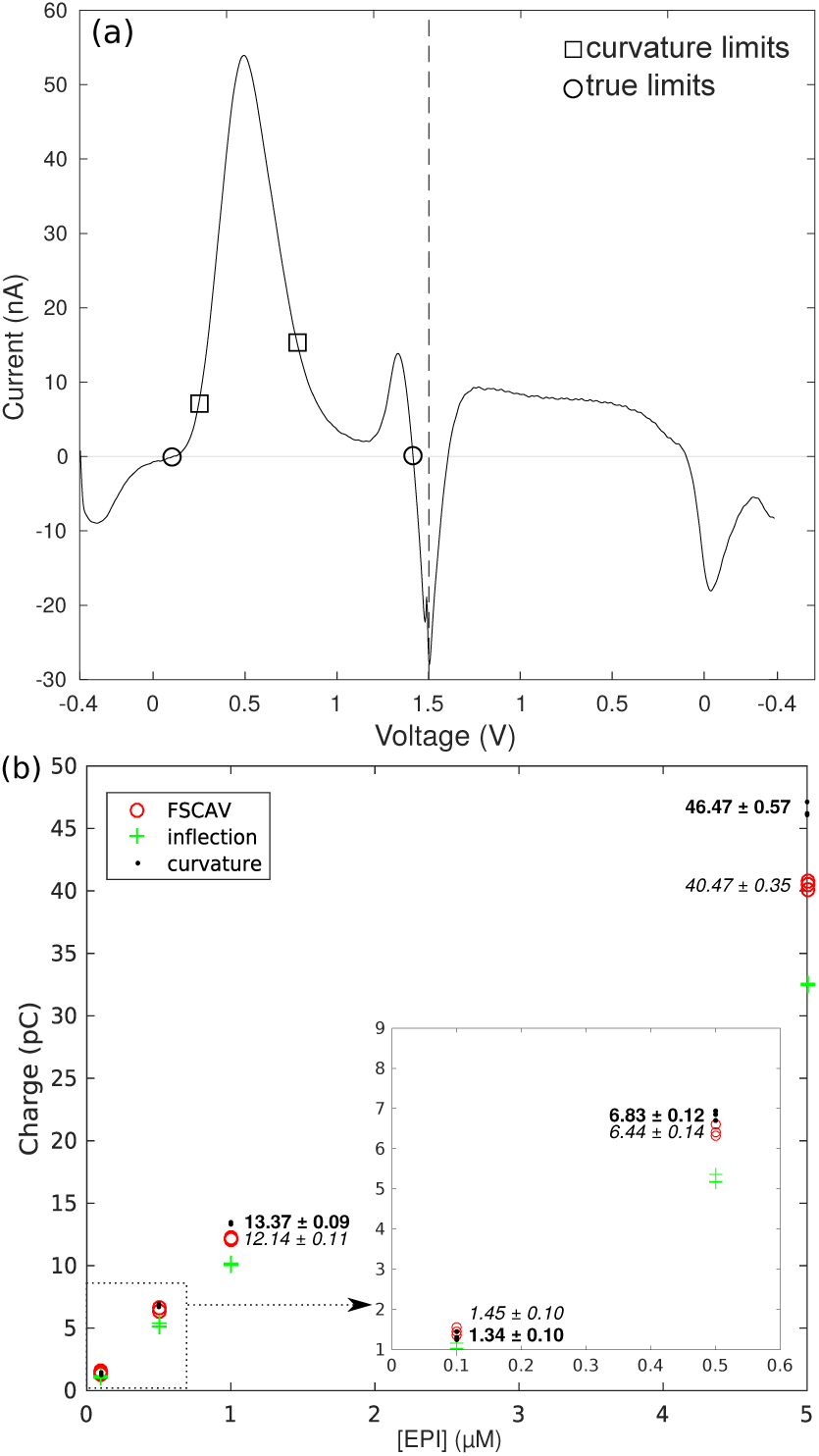
a) Epinephrine voltammogram showing primary and secondary oxidation peaks, super-imposed with curvature and true integration limits. b) Primary-peak oxidation charges for a flow cell experiment with 15 injections of epinephrine at different concentrations, obtained with different pairs of integration limits. Averages and standard deviations for charges using FSCAV (in italic) and curvature (bold) integration boundaries are also shown.

### 3.2 Charge quantification of in vivo measurements

Rapid changes in the brain electrochemistry can lead to faster background current drift [2, 20, 21], which distorts the voltammograms (an example is shown in figure 5 a). The use of true charges as a benchmark for charge quantification depends on the stability of the background current measured with FSCV. Thus, faster in-vivo background-current change can pose a problem for the use of true boundaries with in-vivo data, as we explain below.

**Figure 5:**
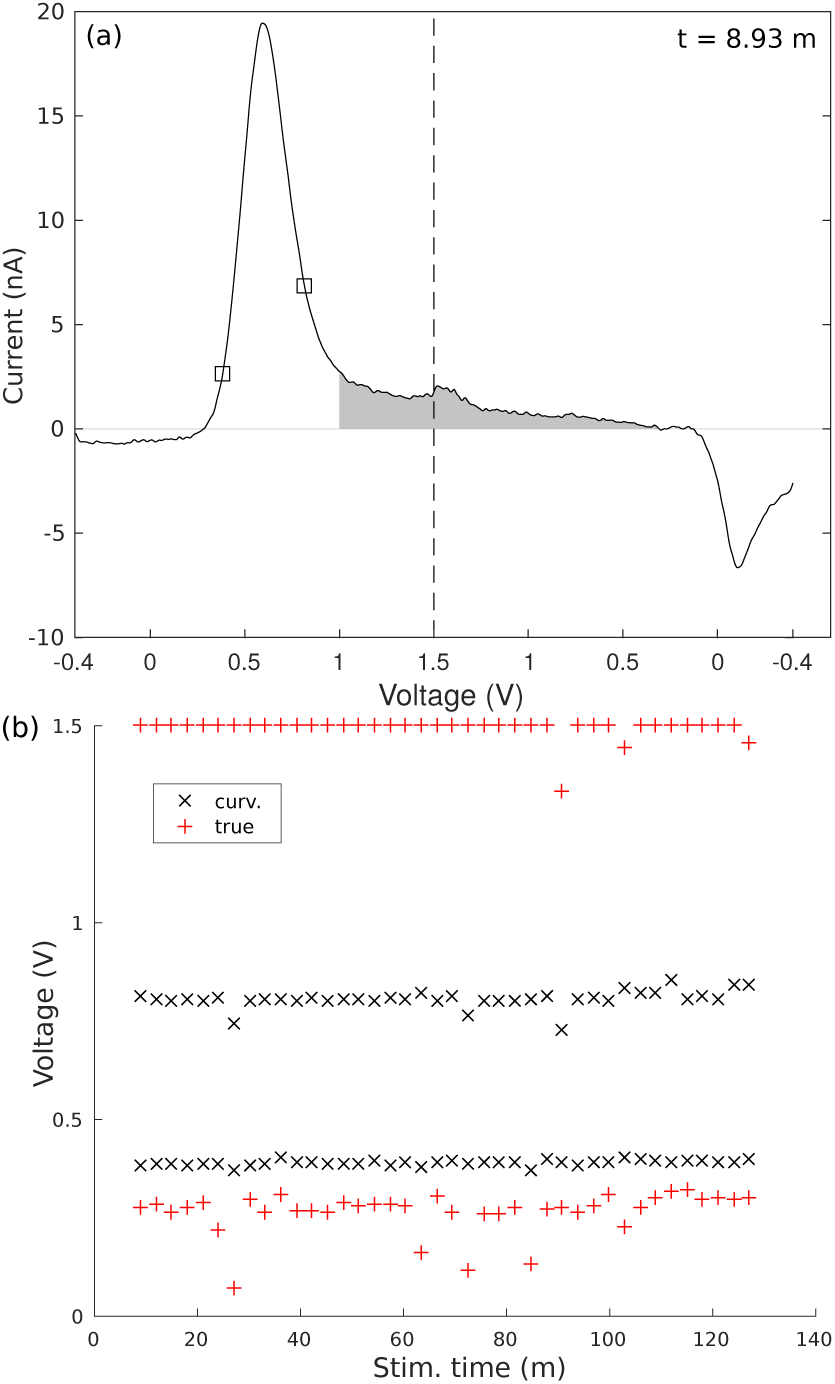
a) Background subtracted voltammogram taken from a rat DBS measurement, superimposed with squares showing the curvature boundaries for the voltammogram. Charge in the shaded region is likely the result of background current drift. b) True and curvature boundaries obtained from background subtracted voltammograms of the entire rat DBS dataset.

Figure 5(a) shows a stimulation-evoked background-subtracted voltammogram from a rodent striatum measurement, superimposed with curvature boundaries. A region of positive, nearly constant current between 1 and 1.5 volts that persists far into the *cathodic* sweep, is likely the consequence of background drift. The shaded region adds a positive bias to the charge computed with true boundaries. Furthermore, a similar behavior is observed in most voltammograms of the data set of Figure 5. As a consequence, in panel (b), which shows true and curvature boundaries for the entire dataset, the right true boundary *x*_*r*_ has been set to the switching potential for most cases.

We contrast the behavior of true boundaries for the experimental data set of figure 5(b) with that of the curvature boundaries, which despite the random variation of the DBS stimulation parameters of the experiment, show little variability (*x*_*l*_ = 0.388 ± 0.007 V and *x*_*r*_ = 0.807 ± 0.022 V), demonstrating the robustness of our charge quantification algorithms.

This robust behavior can be utilized for outlier detection purposes. Values that deviate the most from the average right curvature boundary of the voltammograms in (*x*_*r*_ = 0.807 ± 0.022 V) Figure 5(a) indicate that stimulation artifacts have altered the shape of the voltammograms and the corresponding curvature boundaries have adapted to the shape of each curve 6(a). Here, details of the oxidation-hump region of three voltammograms are shown with their corresponding curvature boundaries indicated by asterisks.

In Figure 6(b) we present the oxidation charges corresponding to the in-vivo experimental data set of Figure 5, as function of the stimulation duty cycle. This panel shows how the response measured by the CFM increases as a function of the increasing stimulation duty cycle.

**Figure 6:**
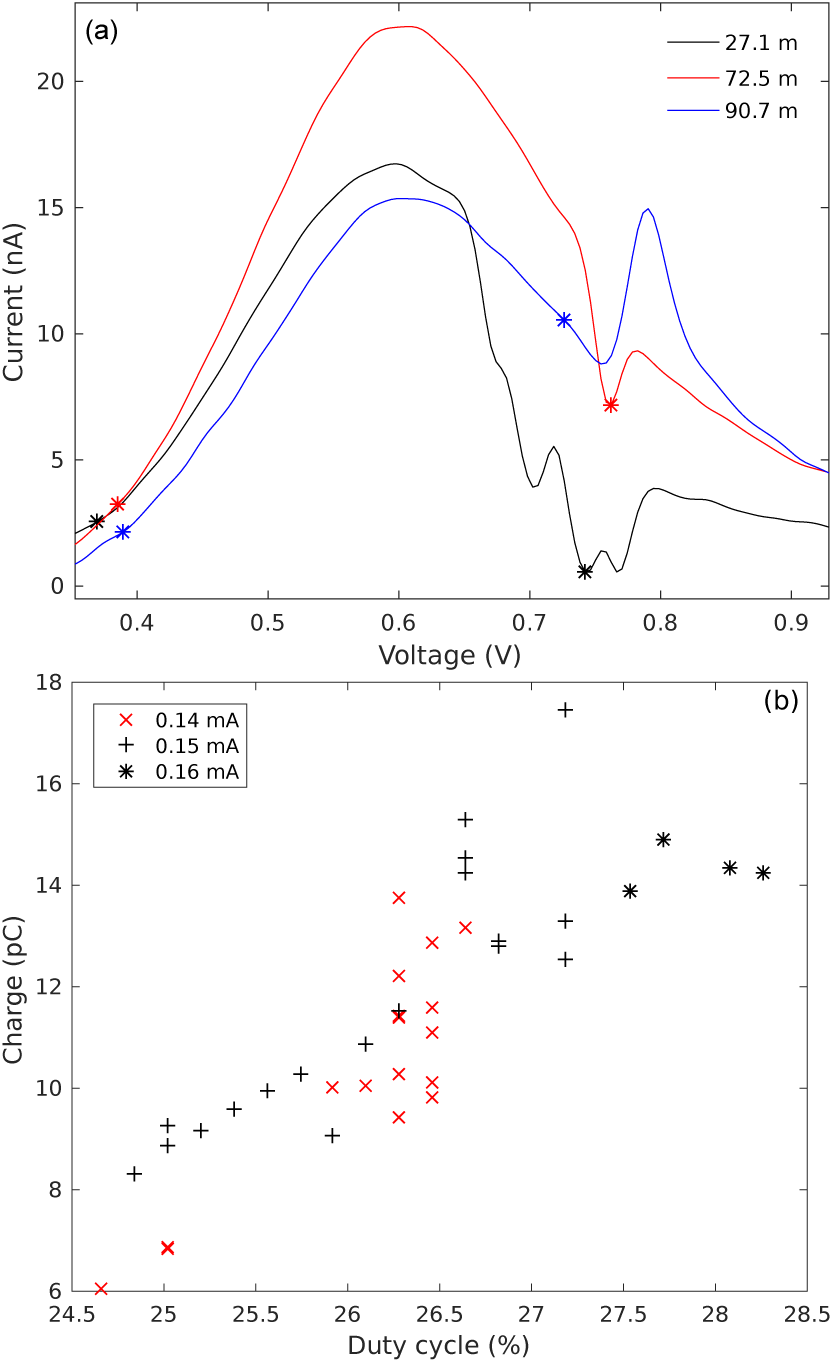
a) Voltammogram regions enclosing the oxidation peak of selected background subtracted voltammograms taken from the data set of Figure 5, superimposed with asterisks showing the curvature boundaries for each voltammogram. (b) Oxidation charges corresponding to the data set of Figure 5, obtained with curvature limits, plotted as function of the duty cycle for the 90 Hz, two-second stimulations.

### 3.3 Quantifying charge produced in reduction reactions

As described in section 2.3, our methods can be used to quantify charged due to reduction reactions, and our algorithms require minimal alterations to do so (finding minima instead of maxima, etc). Figure 7(a) shows the curvature limits for the reduction dips measured in the data set of Figure 3(a). Figure 7(b) we show the corresponding charges obtained with curvature limits, as well as with inflection limits. We note that for reduction reactions, very often a right true limit is not existent (see Figure 8(a) for an example). In consequence charges obtained with true limits are not shown.

**Figure 7:**
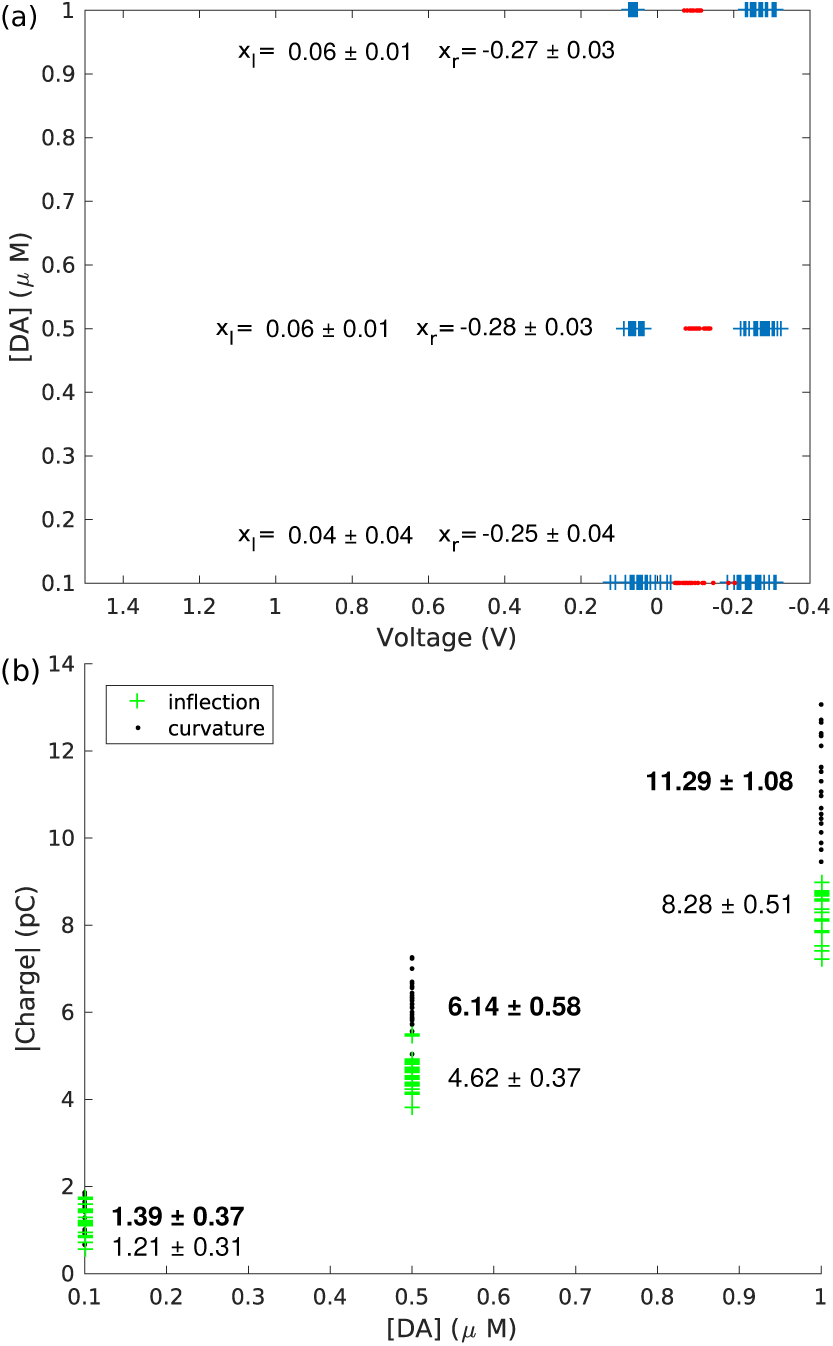
a) Curvature integration limits for the reduction dips of the data set of Figure 3(a). Minimum reduction currents are shown with asterisks. b) Reduction charges (absolute value) for the data set of Figure 3(a), obtained with inflection and curvature (bold) integration boundaries.

**Figure 8:**
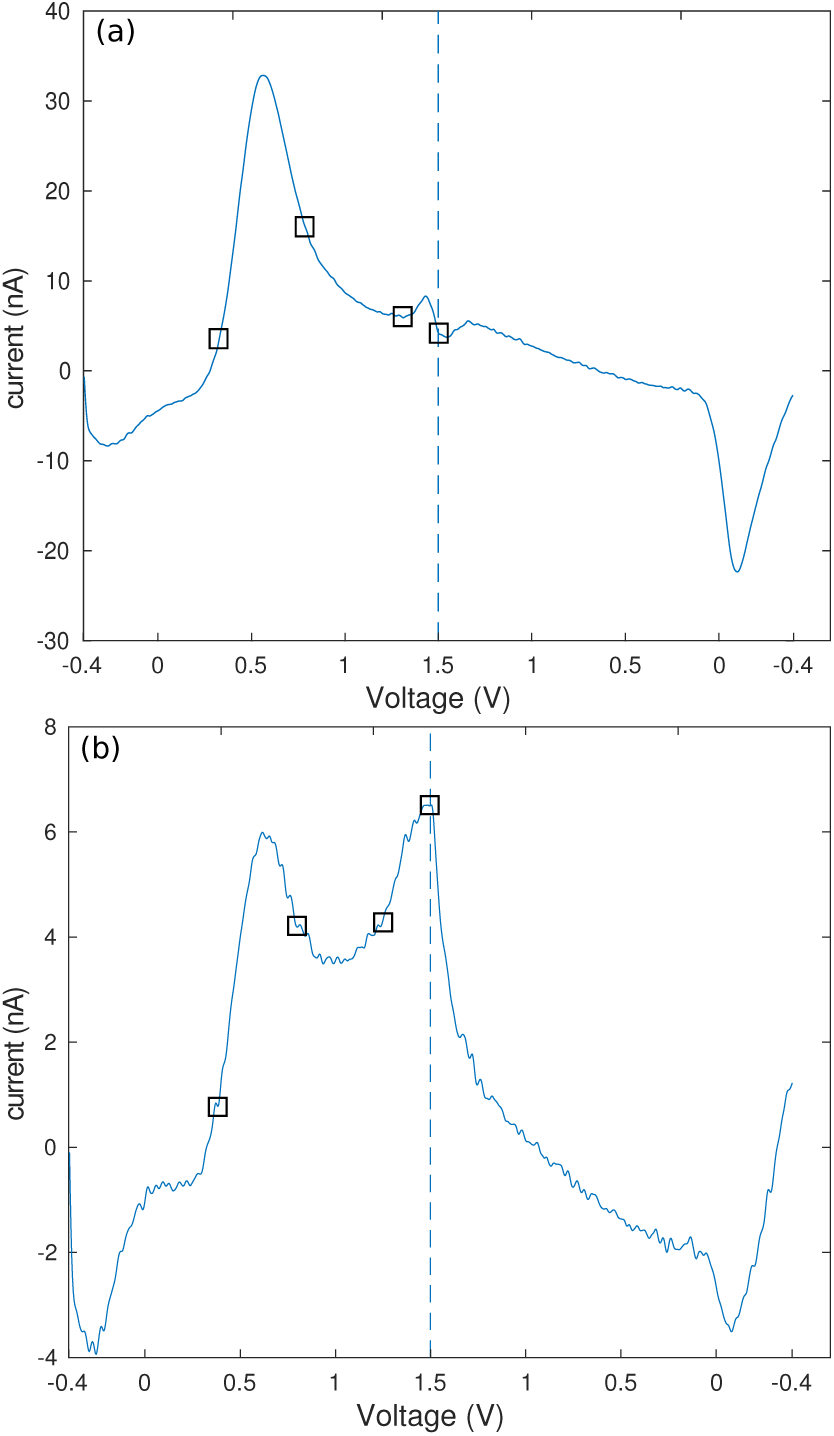
a) Background subtracted voltammogram showing two oxidation peaks due to 1 *µ*M dopamine plus 1 *µ*M adenosine, along with the corresponding curvature boundaries for each peak: 0.33 V and 0.79 V for DA, and 1.31 V and 1.5 V for ADO. b) Background subtracted voltammogram showing two oxidation peaks due to 0.5 *µ*M epinephrine, and the corresponding curvature boundaries for each peak: 0.38 V and 0.80 V for first peak, and 1.25 V and 1.5 V for the second. Vertical dashed lines indicate the switching potentials.

### 3.4 Quantifying charge in voltammograms with multiple oxidation peaks

If multiple electroactive analytes are present in a FSCV measurement, quantifying oxidation due to different species can be challenging. In this section we illustrate how our quantification techniques can aid in that task, when used with cyclic voltammograms displaying multiple humps.

Indeed, the definitions of curvature limits and inflection limits can be used for individual oxidation humps, as we show in Figure 8, where two examples of multiple-oxidation-hump voltammograms, as well as curvature boundaries for each peak are displayed. Notice that in Figure 8(a) the two peaks appear because of two different species, while the voltammogram of Figure 8(b) is the result of the oxidation of epinephrine (see also Figure 4(a)).

## 4 Conclusions

Here, we propose novel methods to quantify charge from oxidation/reduction reactions observed in voltammetric measurements electroactive neurochemicals. While our proposed methods were applied to a dataset collected with a traditional FSCV waveform, this approach can be generalized to complementary voltammetric waveforms dependent on charge quantification, including FSCAV. However, additional studies must be completed to characterize behavior of our approach with novel voltammetric waveforms. Although a definitive selection of integration boundaries is confounded by interferents, background drift, low signal to noise ratio, and other effects, our results show that our quantification methods are equally accurate as competing methods for quantification of dopamine oxidation charge, and are more accurate when applied to other chatecolamines. Additionally, unlike existing charge quantification techniques, our methods can be used to quantify reduction reactions as well as single oxidation or reduction peaks when multiple analytes are present in a sample.

Here, charge analysis has been performed automatically, improving reproducilibity and demonstrating the feasibility of developing automated routines for charge quantification of multiple analytes.

## References

[1] R. M. Wightman, C. Amatorh, R. C. Engstrom, P. D. Hale, E. W. Kristensen, W. G. Kuhr, and L. J. May, “Real-time characterization of dopamine overflow and uptake in the rat striatum,” Neuroscience, vol. 25, no. 2, pp. 513–523, 1988.

[2] D. L. Robinson, A. Hermans, A. T. Seipel, and R. M. Wightman, “Monitoring rapid chemical communication in the brain,” Chemical Reviews, vol. 108, no. 7, pp. 2554–2584, 2008.

[3] M. L. Heien, M. A. Johnson, and R. M. Wightman, “Resolving neurotransmitters detected by fast-scan cyclic voltammetry,” Analytical Chemistry, vol. 76, no. 19, pp. 5697–5704, 2004.

[4] D. L. Robinson, B. J. Venton, M. L. Heien, and R. M. Wightman, “Detecting subsecond dopamine release with fast-scan cyclic voltammetry in vivo,” Clinical Chemistry, vol. 49, no. 10, pp. 1763–1773, 2003.

[5] D. O. Wipf, A. C. Michael, and R. M. Wightman, “Microdisk electrodes. part ii. fast-scan cyclic voltammetry with very small electrodes,” Journal of Electroanalytical Chemistry, vol. 269, no. 1, pp. 15–25, 1989.

[6] M. J. Logman, E. A. Budygin, R. R. Gainetdinov, and R. Wightman, “Quantitation of in vivo measurements with carbon fiber microelectrodes,” Journal of Neuroscience Methods, vol. 95, no. 2, pp. 95–102, 2000.

[7] D. J. Wiedemann, K. T. Kawagoe, R. T. Kennedy, E. L. Ciolkowski, and R. M. Wightman, “Strategies for low detection limit measurements with cyclic voltammetry.,” Analytical chemistry, vol. 63, no. 24, pp. 2965–70, 1991.

[8] Y. Lu, J. L. Peters, and A. C. Michael, “Direct comparison of the response of voltammetry and microdialysis to electrically evoked release of striatal dopamine,” Journal of neurochemistry, vol. 70, no. 2, pp. 584–593, 1998.

[9] K. T. Kawagoe, P. A. T. Garris, and R. M. Wightman, “ph-dependent processes at nafion-coated carbon-fiber microelectrodes,” Journal of Electroanalytical Chemistry, vol. 359, no. 1-2, pp. 193–207, 1993.

[10] B. P. Jackson, S. M. Dietz, and R. M. Wightman, “Fast-scan cyclic voltammetry of 5-hydroxytryptamine,” Analytical Chemistry, vol. 67, no. 6, pp. 1115–1120, 1995.

[11] C. W. Atcherley, N. D. Laude, K. L. Parent, and M. L. Heien, “Fast-scan controlled-adsorption voltammetry for the quantification of absolute concentrations and adsorption dynamics,” Langmuir, vol. 29, no. 48, pp. 14885–14892, 2013.

[12] C. W. Atcherley, K. M. Wood, K. L. Parent, P. Hashemi, and M. L. Heien, “The coaction of tonic and phasic dopamine dynamics,” Chemical Communications, vol. 51, no. 12, pp. 2235–2238, 2015.

[13] M. H. Burrell, C. W. Atcherley, M. L. Heien, and J. Lipski, “A novel electrochemical approach for prolonged measurement of absolute levels of extracellular dopamine in brain slices,” ACS Chemical Neuroscience, vol. 6, no. 11, pp. 1802–1812, 2015.

[14] Y. Oh, C. Park, D. H. Kim, H. Shin, Y. M. Kang, M. DeWaele, J. Lee, H.-K. Min, C. D. Blaha, K. E. Bennet, I. Y. Kim, K. H. Lee, and D. P. Jang, “Monitoring in vivo changes in tonic extracellular dopamine level by charge-balancing multiple waveform fast-scan cyclic voltammetry,” Analytical Chemistry, vol. 88, no. 22, pp. 10962–10970, 2016.

[15] K. H. Lee, J. L. Lujan, J. K. Trevathan, E. K. Ross, J. J. Bartoletta, H. Park, S. Paek, E. N. Nicolai, J. H. Lee, P. H. Min, C. J. Kimble, C. D. Blaha, and K. E. Bennet, “Wincs harmoni: Closed-loop dynamic neurochemical control of therapeutic interventions,” Nature Scientific Reports, 2017.

[16] P. E. M. Phillips, D. L. Robinson, G. D. Stuber, R. M. Carelli, and R. M. Wightman, “Real-time measurements of phasic changes in extracellular dopamine concentration in freely moving rats by fast-scan cyclic voltammetry,” Methods Mol Med, vol. 79, pp. 443–464, 2003.

[17] A. V. Oppenheim, A. S. Willsky, and S. H. Nawab, Signals and Systems (2Nd Ed.). Upper Saddle River, NJ, USA: Prentice-Hall, Inc., 1996.

[18] J. Park, P. Takmakov, and R. M. Wightman, “In vivo comparison of norepinephrine and dopamine release in rat brain by simultaneous measurements with fast-scan cyclic voltammetry,” Journal of Neurochemistry, vol. 119, no. 5, pp. 932–944, 2011.

[19] M. L. A. V. Heien, A. S. Khan, J. L. Ariansen, J. F. Cheer, P. E. M. Phillips, K. M. Wassum, and R. M. Wightman, “Real-time measurement of dopamine fluctuations after cocaine in the brain of behaving rats.,” Proceedings of the National Academy of Sciences of the United States of America, vol. 102, no. 29, pp. 10023–8, 2005.

[20] D. L. Robinson, B. J. Venton, M. L. Heien, and R. M. Wightman, “Detecting subsecond dopamine release with fast-scan cyclic voltammetry in vivo,” Clinical Chemistry, vol. 49, no. 10, pp. 1763–1773, 2003.

[21] D. P. Jang, I. Kim, S. Y. Chang, H.-K. Min, K. Arora, M. P. Marsh, S. C. Hwang, C. J. Kimble, K. E. Bennet, and K. H. Lee, “Paired pulse voltammetry for differentiating complex analytes.,” The Analyst, vol. 137, no. 6, pp. 1428–35, 2012.

